# Maternal Inflammation in Late Gestation Alters Vaccine-Induced Immune Responses in Adult Murine Offspring

**DOI:** 10.64898/2026.04.23.719749

**Authors:** Casey M. Nichols, Dajana Sabic, Jay J. McQuillan, Joyce M. Koenig

**Affiliations:** Parks College of Engineering, Aviation, and Technology; Saint Louis University, St. Louis, MO, USA; Department of Pathology, Microbiology and Immunology, Vanderbilt University Medical Center, Nashville, TN, USA; Department of Pediatrics; Saint Louis University, St. Louis, MO, USA; Department of Pediatrics, Texas Children’s Hospital, Baylor College of Medicine, Houston, TX, USA; Department of Molecular Microbiology & Immunology; Saint Louis University, St. Louis, MO, USA

**Keywords:** maternal inflammation, postnatal immunity, vaccine, immune challenge

## Abstract

**Background:** Intrauterine inflammation, commonly presenting as chorioamnionitis, is variably linked to preterm birth, neonatal infections and postnatal chronic inflammatory disorders. However, the effects of systemic maternal inflammation on exposed fetuses and offspring are less clear. We previously reported inflammatory responses in murine pups born after brief gestational exposure to experimental maternal inflammation. These findings led us to hypothesize that fetal exposure to maternal inflammation could lead to persistent alterations in postnatal immunity.

**Objective:** To test our hypothesis, we examined immune responses to vaccination, a useful measure of immune status, in young adult offspring with late gestational exposure to maternal LPS.

**Design/Methods:** Late-gestation pregnant dams were treated with LPS or saline. Offspring (LPS-exposed or saline controls) were either immunized with the Tdap vaccine or remained unimmunized (naive mice), and were subsequently infected with *Bordetella pertussi*s. Lung and spleen immune responses were assessed by multi-parameter flow cytometry, protein microarray and RT-PCR.

**Results:** We observed that young adult (7 week old) mice exposed to maternal LPS during gestation, vaccinated with TDaP, and subsequently infected with pertussis exhibited lower lung neutrophil but higher CD4+ lymphocyte proportions relative to unexposed controls. In splenic studies, LPS-exposed mice had lower frequencies of CD4+IFNγ+ (Th1) and CD4+IL-17+ (Th17) cell populations. *In vitro* studies of post-vaccination responses to heat-killed *B. pertussis* showed variable levels of IL-2 and IL-4 in splenic cultures from LPS-exposed *vs.* control mice. Vaccinated, LPS-exposed mice showed variable splenic *Stat3* and *NFkb* gene expression levels relative to those of naive LPS-exposed mice.

**Conclusion:** Our present murine studies show that experimental maternal inflammation during late gestation can alter immune response patterns to secondary challenge in young adult offspring. However, whether such intrauterine inflammatory exposure might also influence protective immune function remains to be determined. Our findings lead us to speculate that fetal exposure to systemic maternal inflammation in humans could have long-term implications for protective immunity.

## INTRODUCTION

Alterations of the fetal environment can result in adverse long-term health consequences, consistent with the developmental origins of disease concept proposed by Barker ^1^. Supportive of this, early-life exposure to inflammation has been shown to modulate postnatal innate and adaptive immune development ^2–4^. This influence may be particularly potent when this exposure occurs during the fetal period, whether as a result of placental inflammation ^5^ or in the context of maternal immune activation ^3,6^. Preterm infants born with evidence of fetal inflammation are highly susceptible to sepsis and are at greater risk for childhood asthma ^7,8^, which suggests an adverse effect on developmental immune programming (reviewed in ^9,10^). In addition, some evidence indicates a link between fetal inflammatory exposure and immune-related neurologic disease in adulthood ^11^. However, the processes by which inflammatory exposure at critical developmental windows may influence protective immunity in later life have not been fully defined. Experimentally-induced antenatal inflammation can lead to exaggerated inflammatory responses in exposed neonatal offspring (reviewed in ^6,9^) and ^12^ through processes that remain incompletely defined.

Vaccine-induced responses are key metrics of immune integrity ^13^. Notably, human preterm infants exhibit variable protective responses to vaccines ^14^. In a study of pertussis immunization of extremely premature neonates, a protective immune response was observed in less than half of recipients ^15^, although whether exposure to antenatal inflammation might have been an underlying factor was not specifically addressed. Altered fetal and postnatal immune responses in the context of maternal and intrauterine inflammation have been reported in animals and humans ^16–19^. These observations led us to question whether fetal inflammatory exposure might adversely influence protective immune responses in affected offspring.

To test this hypothesis, we examined vaccine responsiveness as a determinant of immune status^13^. Immune responses to pertussis vaccination and *B. pertussis* infection have been well characterized in several murine models ^20–22^. We incorporated various aspects of these models into our existing one of antenatal inflammation ^12^ to determine the effects of maternal inflammation on vaccine-related immune response patterns in LPS-exposed offspring and saline treated controls.

## METHODS

### Study design

Timed-pregnant C57Bl/6J mice (Jackson Labs, Bar Harbor, ME) were housed in a pathogen-free biohazard barrier facility in microisolator cages, and received a standard diet. To induce maternal inflammation, lipopolysaccharide (LPS, *Escherichia coli* 0111:B4, Sigma, St. Louis MO) diluted with injection-grade saline (USP 0.9% NaCl, Sigma) was administered IP (50-75 μg/kg) to pregnant dams (E17); control dams received equivalent volumes of the sterile saline vehicle, as we previously described ^12^. Dams were allowed to deliver spontaneously; day of delivery was considered to be D0. We next assessed immune status in LPS-exposed or control offspring by examining their responses to immune challenge (**Fig. 1**). Briefly, offspring in each group received a ‘priming’ immunization (or the same volume of sterile saline for naive animals) via the *i.p.* route on postnatal day (PND) 5. All mice (LPS-exposed or unexposed controls) subsequently received a second treatment (‘booster’ vaccine (vax) or saline for naïve mice) 4 weeks after the first injection. To stimulate an immune response, some mice received a non-lethal intranasal inoculation of *B. pertussis* (2.5 x 10^6^ CFU) two weeks after the booster injection, as reported in previous studies ^21,22^. On post-infection day (PID) 7 (or 7 days after sham infection), whole blood and solid organs were collected from offspring of LPS-treated (LPS-exposed) or saline-treated (control) dams.

**Figure 1.**
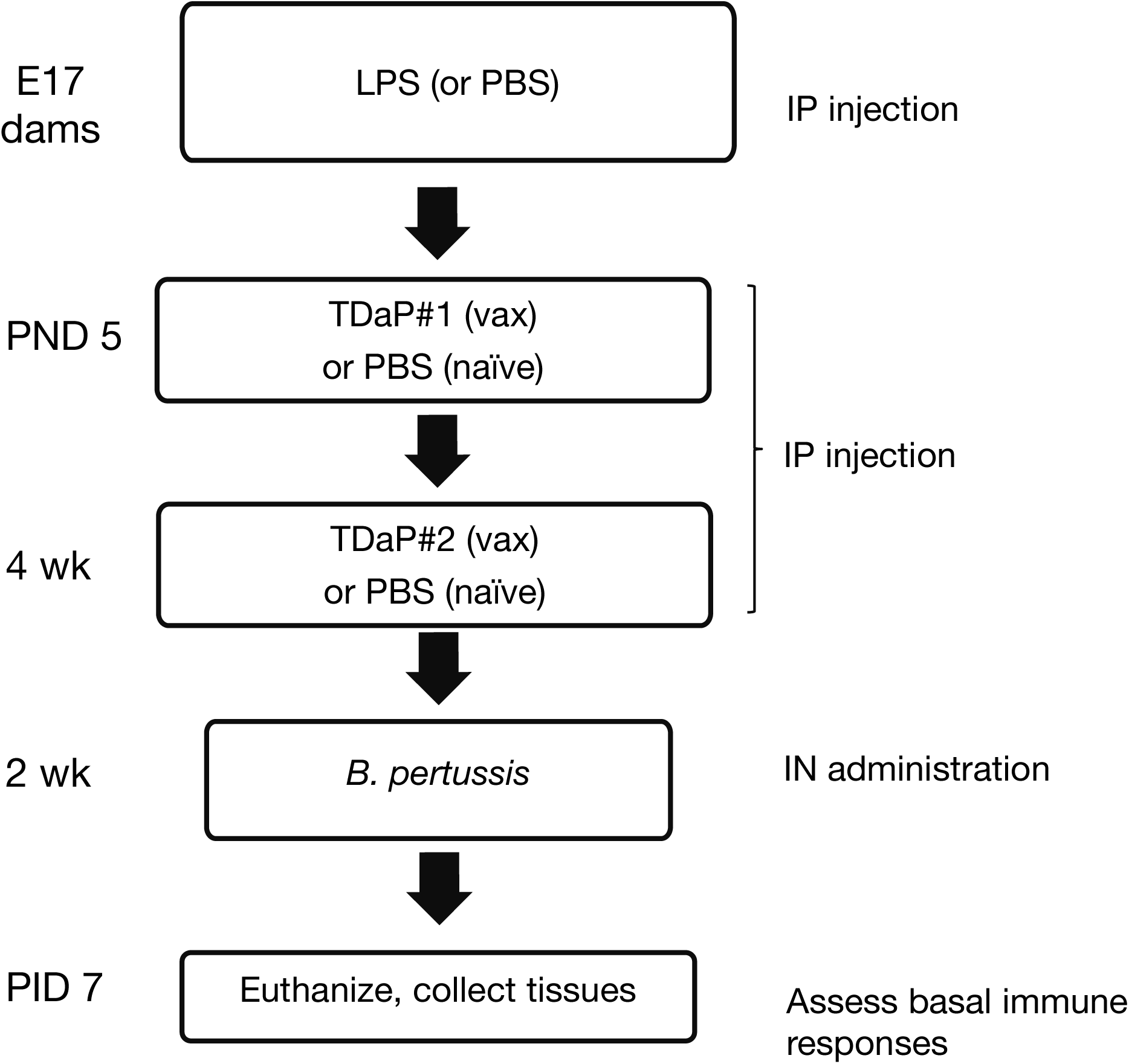
Study design. Timed-pregnant (E15) dams received intraperitoneal (IP) LPS (or a similar volume of endotoxin-free PBS for controls). Spontaneously delivered pups received IP TDaP (or PBS for controls) on postnatal day (PND) 1. A booster IP dose of TDaP was given to vaccinated offspring (or PBS to controls) 4 weeks later. Vaccinated or naïve (unvaccinated) control mice were infected with *B. pertussis* via the intranasal (IN) route 2 weeks after the booster vaccine, and euthanized on post-infection day (PID) 7.

### Blood and tissue harvest

Whole blood was obtained via cardiac puncture and solid organs (lungs, spleen) were harvested from euthanized mice, as per an approved IACUC protocol. Harvested organs were processed to yield single cell suspensions, as described ^12^. Lungs underwent additional digestion in a solution of IMDM containing hyaluronidase, collagenase I and DNase I ^23^. Single cell suspensions derived from spleens and lungs were stained for flow cytometric analysis. To quantify viable myeloid and lymphoid cell subsets, cell suspensions were stained with specific fluorochrome-labeled mAb. To identify T-helper cell subsets, harvested cells were processed by intracellular cytokine staining (ICS). Pertussis-specific responses were determined in splenocytes stimulated for 7 d with heat-inactivated *B. pertussis*, as described ^24,25^.

### Vaccine

For these studies we utilized the Tdap vaccine (Boostrix®, GlaxoSmithKline, East Durham, NY), a tricomponent acellular pertussis vaccine (pertussis toxoid (PT), 8 μg; pertactin, 2.5 μg; filamentous hemagglutinin (FHA), 8 μg) formulated with diphtheria toxoid (2 IU) and tetanus toxoid (20 IU). The undiluted vaccine was administered to experimental mice at a volume of 0.1 mL (20% of the human dose) via the *i.p*. route ^22^.

### Bacteria and bacterial antigens

The wild-type strain of *B. pertussis* (BP536, kind gift of Dr. Stephen Barenkamp) ^26^ was used for the intranasal challenge study. *B. pertussis* was grown on Bordet-Gengou agar plates supplemented with 15% sheep blood (BGA) for 24 h (37°C), followed by culture in Stainer-Scholte liquid medium for 18 h to achieve the exponential phase ^26^. Heat-inactivated bacteria were utilized for *in vitro* determinations of pertussis-specific immune responses (described below).

### Antibodies and reagents

Fluorochrome-labeled antibodies were utilized for cell surface staining: L/D Aqua (viability stain), CD45-, CD3e-Cy5, CD4-PE, CD8a-V450, CD45-APC, CD11b-AF700, CD11c-PECy7, Gr-1-PerCP-Cy5.5, Ly-6G-PE-CF594, MHCII-FITC. For intracellular staining (ICS), antibodies also included: IL-4-PerCPCy5.5, IL-10-BV711, IL-17-PE, INFy-BV786, CD8a-V450. Specific IgG subset controls were included as indicated. All antibodies were purchased from Becton-Dickinson (Franklin Lakes, NJ).

### Flow cytometric analyses

For surface staining, spleen or lung single cell (10^6^) suspensions were incubated with mAb, as described ^12^. For intracellular cytokine detection in spleens or lungs, cell suspensions were first stimulated with phorbol myristic acid (PMA, 50 μg/ml), ionomycin (1 μg/ml) and brefeldin A (5 μg/ml) for 4 h at 37°C (unstimulated samples were incubated with IMDM/2% fetal bovine serum). After the initial surface antigen staining process, cells were fixed and permeabilized, then stained with a cocktail of anti-cytokine mAb. Cells were acquired by a multi-parameter flow cytometer (BD LS-II) within 24 h of staining in order to preserve the fluorescent signal. Acquired data were analyzed in viable cells using the FlowJo software program (Tree Star, Ashland OR). After gating for the viable CD45+ leukocytes, myeloid and lymphoid cell populations were identified using forward- and side-scatter characteristics and specific surface markers with the guidance of IgG subtype-specific Ab controls. Identification of myeloid cells was based on surface expression of Gr-1 or Ly6G and CD11b. CD3+ lymphoid cell subsets were identified by their surface expression of CD4 or CD8. Identification of T helper (Th) cell populations within CD4 subsets was based on the intracellular expression of IFNγ (Th1), IL-4 (Th2) or IL-17 (Th17).

### Pertussis-specific splenic cytokine responses

To assess the expression levels of pertussis-specific cytokine responses, splenocytes (10^6^ cells) from vaccinated LPS-exposed or control mice were cultured in media only (IMDM/2% fetal bovine serum; negative control) or in the presence of heat-inactivated whole cell *B. pertussis* (10^7^/mL) as an immune challenge ^20,22^. After 6 days of culture, supernatants were aliquoted and stored at -80° C until batch analysis for the detection of immune cytokines, as described below.

### Pertussis-specific cytokine protein array

Supernatants of spleen cells incubated with heat-inactivated *B. pertussis* bacteria were tested for their content of immune cytokines (IFN-γ, IL-1β, IL-2, IL-4, IL-6, IL-17A, IL-10, TNF-α) using a protein microarray (Cytometric Bead Array, BD Mouse Th1/Th2/Th17 CBA Kit, Cat #560485). Cytokine capture beads for the various cytokines were prepared as per the manufacturer’s instructions. Cytokine bead suspensions (50 µL) were combined with culture supernatants or standards (50 µL) in each well of a 96-well plate. The plate was incubated at RT in the dark for 2 h, then washed twice with wash buffer, as per manufacturer’s instructions. Beads were resuspended in 300 µL of wash buffer and acquired by a multiparameter flow cytometer.

### RNA extraction

Freshly harvested spleen and lung tissue samples, or post-culture cell pellets, were stored in RNAlater^TM^ (Ambion,Inc USA) at 4°C until further processing. Total RNA was extracted using the RNeasy^TM^ mini kit (Qiagen, Hilden, Germany) according to the manufacturer’s instructions and as described ^27^. Briefly, cells were centrifuged and harvested, supernatants were removed and RNA extracted from each cell pellet and resuspended in lysis buffer supplemented with β-mercaptoethanol to disrupt the cells. Cells were lysed and each lysate was homogenized and mixed with equal volume of 70% ethanol. Cell lysates were transferred onto RNeasy mini-spin columns and DNA was removed using DNase digestion/ treatment using RNase-Free DNase Set (Qiagen, Hilden, Germany.) The RNA Integrity Number (RIN) of all RNA samples were measured using an Agilent 2100 Bioanalyzer and Agilent RNA 6000 nano kit (Agilent Technologies, Santa Clara, CA, USA); with a minimum RIN of 7.5 used as the criterion for inclusion in gene expression analysis. The concentration of each individual RNA sample was measured using a Qubit RNA BR Assay (Invitrogen) in triplicate. Several blood samples were collected for PBMC isolation, treatment and extraction of RNA and only RNA samples with the highest RIN numbers (all above 8) were included for PCR.

### qPCR

Reverse transcription was performed by adding 4.5 µL iScript Reverse Transcription Supermix (Bio-Rad Cat#1708840) to the 15.5ul RNA template (10 µg/ml). The reaction was incubated at 25° C for 5 min, 42° C for 30 min, and 85° C for 5 min, and then stored at −20°C. denaturation at 95°C for 15 s followed by an annealing/amplification step at 58°C for 4 min. Reactions were stored at −20°C. For expression profiling using qPCR each qPCR reaction contained 2 µL of diluted template, 1x SsoAdvanced universal SYBR supermix (Bio-Rad) and 1x PrimePCR assays (Bio-Rad). Reactions were performed in 3 technical replicates at 20 µL final volume. Real-time PCR was performed using the CFX96 real-time PCR systems (Bio-Rad): activation at 95° C for 2 min, 40 cycles of denaturation at 95° C for 5 s and annealing/elongation at 60° C for 30 s. Specificity of target amplification was confirmed by melting-curve analysis. The following genes were evaluated: *Stat3*, 5-’GGGCCTGGTGTGAACTACTC-3’; *Nfkb*, 5-’CCGTCTGTCTGCTCTCTCT-3’. Βeta-actin was used as a control. Expression data were analyzed using the CFX manager 3.1 software based on the ΔΔCq method and graphed in GraphPad Prism 5.02 (GraphPad Software Inc., La Jolla, CA, USA).

### Statistical analyses

Statistical analyses were performed using the GraphPad 7 (Prism) software program. Pairwise comparisons were made using Student’s *t*-test or the nonparametric Mann-Whitney U-test. Multiple group differences were evaluated by ANOVA and Tukey’s *post hoc* test. Results are expressed as means ± SEM. A p value < 0.05 was considered to be statistically significant.

## RESULTS

### Maternal LPS treatment alters myeloid and lymphoid responses to vaccination in offspring

To assess the effects of antenatal LPS or saline exposure on innate responses in offspring following immune challenge, vaccinated LPS-exposed (LPS-Vax) and control (Ctrl-Vax) mice were infected with pertussis. At the expected peak of infection (PID 7), we utilized flow cytometric analysis to determine myeloid cell populations in spleens and lungs ^24^. The proportions of myeloid cells (Gr1+CD11b+, primarily inclusive of neutrophils and other myeloid subsets; and Ly6G+CD11b- monocytes) were analyzed in CD45+ gated populations of cell suspensions. As shown (**Fig. 2A**), in LPS-Vax mice, Gr1+ neutrophil proportions were lower in both the spleens and lungs relative to Ctrl-Vax mice. In contrast, monocyte proportions were suppressed in the spleens but not lungs of LPS-Vax mice (**Fig. 2B**). Similarly, to address adaptive cellular responses, we analyzed CD3+ lymphoid populations in the same cell suspensions. As shown (**Fig. 2C**), CD3+CD4+ proportions were higher in the lung suspensions of LPS-Vax *vs.* Ctrl-Vax mice, while a similar, but non-significant trend was observed in spleens. In contrast, no differences were determined in the proportions of CD3+CD8+ cells between groups in the spleens or lungs.

**Figure 2.**
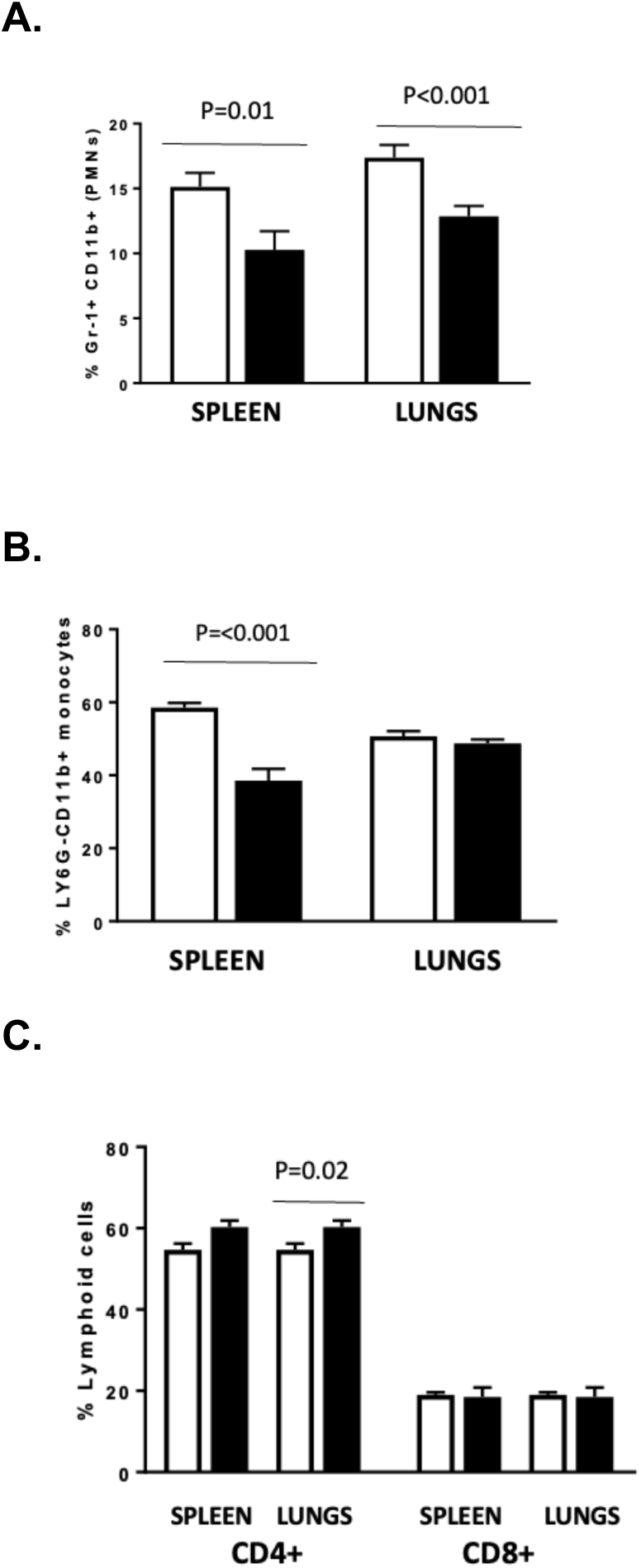
Myeloid responses in the spleens and lungs of vaccinated offspring. Single cell suspensions obtained from vaccinated controls (Ctrl-Vax, *open bars*) or from LPS-exposed offspring (LPS-Vax, *black bars*) were stained for flow cytometric analysis. Neutrophils (**A**., Gr-1+CD11b+), monocytes (**B**., Ly6G-CD11b+ cells), and lymphocyte subsets (**C**., CD4, CD3+CD4+CD8- and CD8, CD3+CD4-CD8+) were analyzed and compared. Data here and in subsequent panels represent the results of one of 3 studies; *n* = 4 mice per group, X ± SEM.

### Maternal LPS suppresses T-helper cell responses in vaccinated offspring

The induction of adaptive immune Th cell responses, particularly those involving Th1 and Th17 cells, is an important aspect of protective cell-mediated immunity to infection ^28^. To examine the effects of antenatal LPS exposure and vaccination on the postnatal induction of specific Th cell subsets, we performed comparative intracellular cytokine analyses of Th cells in the spleens and lungs of offspring. As shown (**Fig. 3A**), LPS-Vax offspring had lower splenic proportions of CD4+IFNγ+ and CD4+IL-17+ cell populations, but higher proportions of CD4+IL-4+ cells, relative to Ctrl-Vax mice. A similarly suppressive pattern was observed for splenic CD8+IFNγ+ cells in LPS-Vax *vs*. Ctrl-Vax mice (**Fig. 3B**), while CD8+IL-17+ cell proportions in LPS-Vax mice also showed a diminished but non-significant trend. No sex-specific differences in spleen cytokine responses were detected between male and female mice in either the LPS-Vax or Ctrl-Vax groups (data not shown), thus data from both sexes were combined for these and following studies.

**Figure 3.**
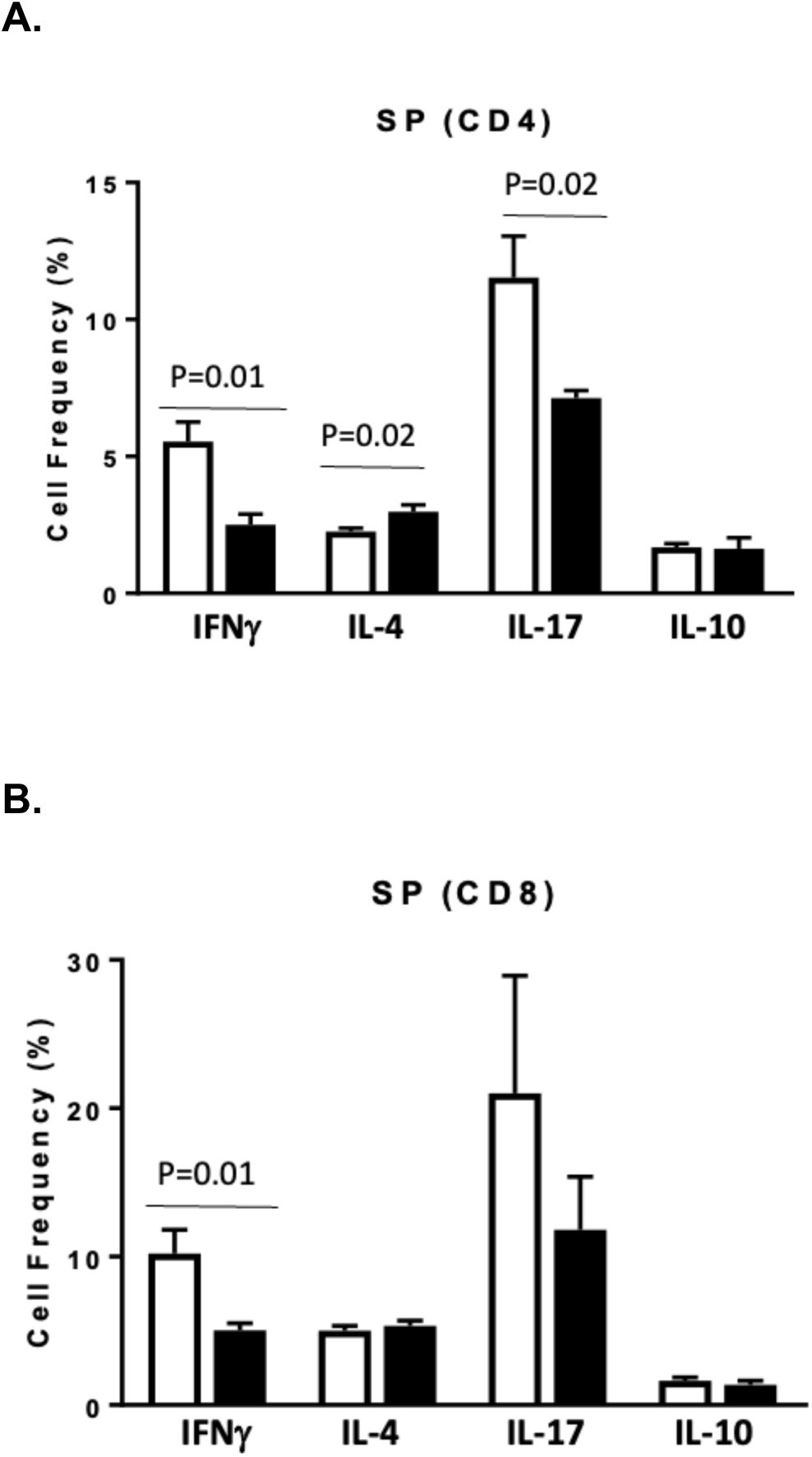
Intracellular cytokine expression patterns in splenic lymphocytes of vaccinated and pertussis-infected offspring. Stimulated single cell suspensions were stained by ICS. Proportions of splenic CD4+ (**A.**) and CD8+ (**B.**) cells, with intracellular expression of various cytokines were compared in cells from Ctrl-Vax (*open bars*) or LPS-Vax (*black bars*) offspring.

### Pertussis antigen-induced release of splenic cytokines

To identify specific cytokine responses to pertussis vaccination, we utilized a flow cytometry-based protein array assay to assess *in vitro* cytokine release induced by exposure to heat-killed pertussis bacteria in spleen cells obtained from mice that were either naïve (not immunized) or that had received vaccine treatment. We determined levels of IL-2 and IL-4 in splenic culture supernatants of mice that varied based on having been vaccinated or remaining naive (**Fig. 4**). Regarding IL-2 levels, in the LPS-exposed group we found lower levels in the vaccinated mice compared to their naïve, unvaccinated counterparts (a pattern not observed in the control offspring group). In addition, IL-2 levels in the vaccinated, LPS-exposed mice were lower than in the vaccinated control offspring. These patterns were observed whether mice were sham infected or had been infected with pertussis (**Fig. 4a).** A different pattern was observed for IL-4 and IL-6. In contrast to IL-2, IL-4 levels were elevated from vaccinated mice compared to their naïve counterparts in both LPS-exposed and control offspring groups, regardless of infection status (**Fig. 4b**). However, as also observed for IL-2, IL-4 levels were lower in the vaccinated LPS-exposed mice relative to vaccinated control offspring for both sham and infected groups. Similarly, for both the LPS-exposed and control offspring, IL-6 levels were higher in vaccinated mice *vs.* their naïve counterparts, but only in the infected groups (**Fig. 4c**). With respect to IL-17 levels, we observed a similar but non-significant trend for infected animals: levels tended to be higher in vaccinated mice in both LPS-exposed and control offspring; however, levels in sham infected groups were nearly undetectable. Levels of the other cytokines assayed in splenic cultures (IFNγ, IL-1β, IL-10, TNFα) were not notably different between naïve and vaccinated groups for either LPS-exposed or control mice (data not shown).

**Figure 4.**
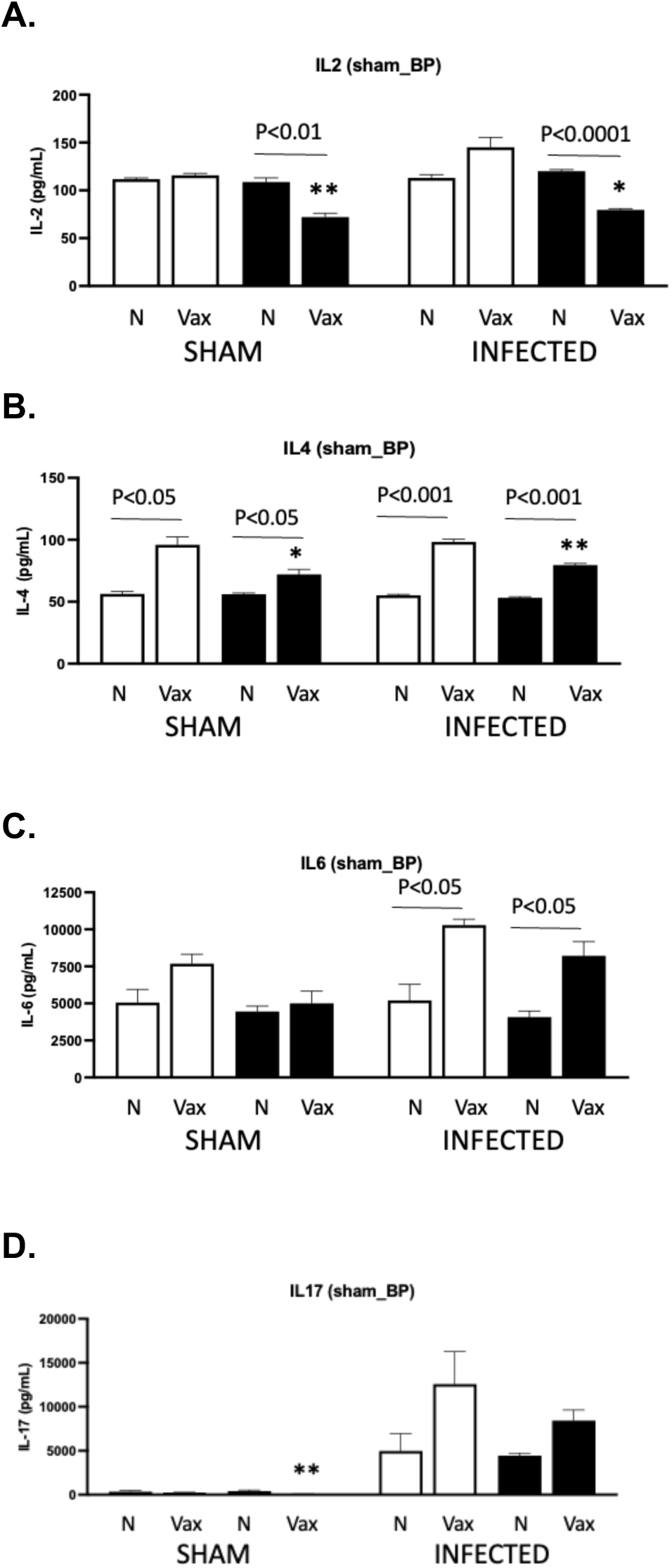
Pertussis-induced cytokine patterns in supernatants of splenocytes from infected naïve (non-vaccinated) or vaccinated offspring. On PID 7 following pertussis infection, spleen cells isolated from mice that either remained naïve (N, non-vaccinated) or that received 2 vaccinations (Vax) were cultured *in vitro* with heat-killed *B. pertussis*. Culture supernatants (D7) were analyzed by protein microarray to assess inflammatory cytokine patterns in spleens of vaccinated offspring of Controls (*open bars*) or LPS-exposed (*black bars*) groups.

### Key inflammatory signaling pathways in LPS-exposed and vaccinated offspring

To begin to understand the mechanisms underlying the suppressed Th1 and Th17 responses observed in LPS-exposed mice, we studied gene expression levels of *Stat3*, the canonical Th17 signaling molecule ^29^ and *Nfkb*, a master regulator of immune and inflammatory responses ^30^. We found lower gene expression levels of *Stat3* in naïve LPS-exposed *vs.* naïve control mice, while there were no differences between vaccinated LPS-exposed and control mice (**Fig. 5**). Additionally, we observed lower *Stat3* gene expression levels in vaccinated LPS-exposed and control mice compared to respective naive mice. Gene expression levels for *Nfkb* were also lower in LPS-vax *vs.* naïve mice, while no differences were observed between Ctrl-vax *vs.* Ctrl-naive mice.

**Figure 5.**
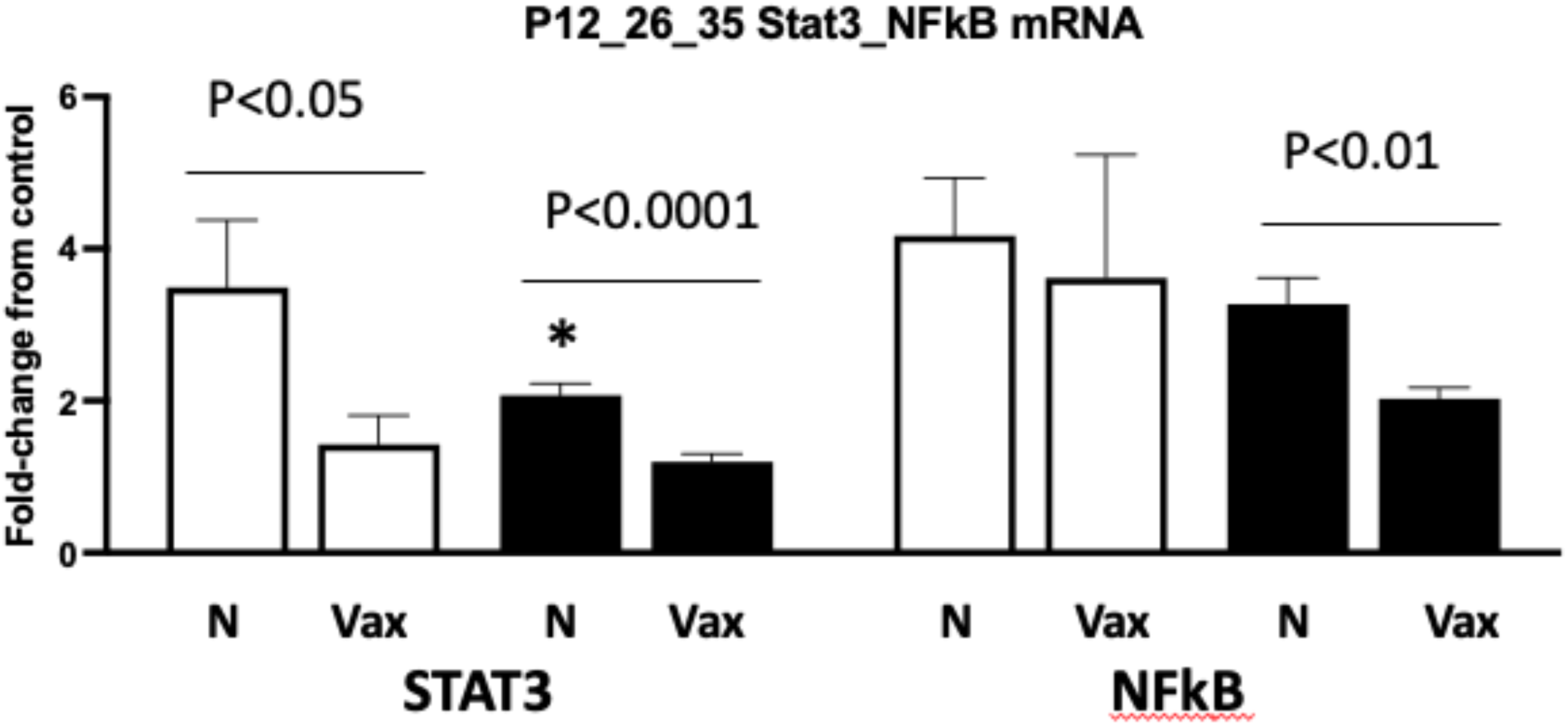
Spleen mRNA expression in infected naïve (unvaccinated) or vaccinated LPS-exposed and control offspring. Data indicate the fold-change in gene expression levels relative to control expression in mRNA extracted from spleens of mice (N: naïve; Vax: vaccinated) that were LPS-exposed (*black bars*) or unexposed controls (*white bars*). Data from 9-20 mice in each group are expressed as X ± SEM. *P <0.05, Ctrl N *vs.* LPS N.

## DISCUSSION

We investigated the effects of maternal inflammation on postnatal immune function by evaluating offspring response to pertussis vaccination and subsequent secondary challenge with non-lethal infection. We now report our findings showing that LPS-induced maternal inflammation can lead to the suppression of certain vaccine-induced immune responses in young adult offspring.

Central to these findings are the immunomodulatory roles of the T-helper cell subsets Th1, Th2, and Th17. The Th1 subset is characterized by IFN-γ production and is important for host defense against intracellular pathogens. Th2 cells produce IL-4, IL-5, and IL-13 and are implicated in allergy and defense against parasites. Pertinently, neonatal immunity in humans has been characterized by a suppressed Th1 response and a pro-Th2 bias that may limit protective immunity ^31,32^. Th17 cells produce IL-17 and have an important role in autoimmunity and extracellular pathogen defense ^33^. The Th1 and Th17 subsets are critical for host defense against pertussis, supporting both clearance of the initial infection and long-term immunity. Production of IFN-γ by Th1 cells is crucial for the recruitment and activation of myeloid cells ^34^, while Th17-mediated release of IL-17 during pertussis infection is necessary to attract neutrophils that mediate bacterial clearance in the lungs ^35,36^.

We previously reported enhanced basal Th1 and Th17 responses in offspring born to LPS-treated dams in our model of antenatal inflammation ^12^. Similar observations have been made in human preterm infants born after chorioamnionitis or following sepsis ^37–39^. In contrast, in our present studies, we found that vaccination and subsequent infection of LPS-exposed offspring led to suppressed splenic Th1 and Th17 responses, as indicated by respectively reduced proportions of IFN-γ+ and IL-17+ CD4 lymphocytes and IFN-γ+ CD8 lymphocytes. This observation was notable given that the C57Bl/6 mouse strain used in our studies has a known Th1 bias ^40^. In contrast, we found elevated proportions of splenic Th2-type (CD4+IL-4+) lymphocytes in the LPS-exposed offspring, a finding which could reflect the Th2 bias associated with the Pa vaccine ^41^. In studies to assess pertussis-specific responses, we observed that the splenocytes of vaccinated LPS-exposed mice released lower IL-2 and IL-4 levels than those measured in vaccinated control offspring. Taken together, the altered immune response patterns that we observed in our LPS-exposed mice are reminiscent of several observations in human infants exposed to maternal inflammation. For example, in a study of infants exposed to maternal HIV infection but not themselves HIV-infected, immunization was associated with a reduction of *in vitro* T cell functionality in response to various antigens, including pertussis ^42^. Human preterm infants exposed to prenatal inflammation were also shown to exhibit suppression of Th1 function ^43^. Our present findings in young adult mice also suggest that alterations in T cell functionality following fetal exposure to systemic maternal inflammation may have the potential for longevity ^7,43^.

In addition to the alterations detected in adaptive cellular function, we observed differences in innate immune responses of LPS-exposed mice. We determined diminished neutrophil proportions in the spleens and lungs of LPS-exposed following. Neutrophils have been shown to interact with and mediate Th17 cell function in order to amplify anti-microbial actions ^45^. Conversely, Th17 cells release cytokines, such as IL-17, that attract neutrophils and facilitate their recruitment to infectious sites ^44^. Thus, a diminution of neutrophil populations combined with lower percentages of Th17 cells could limit neutrophil-Th17 interactions critical to the resolution of infection ^35,36,44^. Additionally, since neutrophils are important in modulating lymphoid function (reviewed in ^45^), a diminished presence in the spleens of LPS-exposed offspring could potentially impact adaptive immune responses ^46^. Pertinently, suppressed innate immune responses (‘tolerance’) to a secondary stimulus has been reported in myeloid cells of fetal sheep following intrauterine exposure to inflammation ^16,47^. Immune tolerance induced by LPS has also been associated with diminished neutrophil infiltration in respiratory tissues and in the brain in several rodent studies ^48,49^. These observations suggest that the diminished myeloid proportions that we observed in the spleens of vaccinated LPS-exposed mice could be the result of inflammation-related tolerance mechanisms.

In initial studies to examine key signaling pathways, we found that vaccination was associated with a lower splenocyte expression of the inflammatory gene *Stat3* (the canonical regulator of Th17 differentiation ^29^) in naïve LPS-exposed offspring. Expression levels were lower for vaccinated mice in both control and LPS-exposed offspring. Expression levels of *Nfkb* (a master regulator of immune and inflammatory responses) were also lower in vaccinated LPS-exposed but not in control offspring. Similar immunosuppressive phenotypes have been observed in fetal sheep and human infants following gestations with chronic maternal infection ^47,50^. While LPS classically upregulates Stat3 and Nfkb via TLR-4 ^51^, there is evidence that Stat3 function is time-dependent in LPS signaling ^52^, potentially explaining this paradoxical suppression. While we did not specifically examine the activation status of these genes, limitations in cytokine synthesis mediated by STAT3 and NF-κB could explain the lower expression of the Th17 and Th1 cytokines ^53^ that we observed. Additional investigations of signaling pathways associated with STAT3 and NF-κB that mediate the dysregulated Th1 and Th17 responses associated with other disorders (reviewed in ^54^) could extend our mechanistic understanding of how inflammatory exposure might promote an immunosuppressive phenotype.

Developmental immaturity is an important barrier to the optimization of vaccine-induced protective immunity in preterm infants. The suppressed Th1 response and pro-Th2 bias reported in neonates ^31,32^ contrasts with the Th1 dominance seen in older children and adults. These differences may weaken vaccine efficacy and increase infection risk ^55^. Vermeulen *et al* found that nearly half of extremely preterm infants failed to exhibit a protective Th1 response to pertussis immunization, while others showed transient responses that waned after one year ^15,56^. Our present studies suggest that fetal exposure to inflammation could have an added negative impact on postnatal immune status. It is tempting to speculate that biologically immature human preterm infants, already hampered by suboptimal immune responses ^19^, could experience added suppression of vaccine-induced immunity following inflammatory exposure during gestation.

In conclusion, our murine studies show that late-gestation fetal exposure to maternal inflammation can alter postnatal vaccine-induced immune response patterns in offspring. Neonatal mice are a useful model of human immune responses to pertussis immunization and to *B. pertussis* infection (^22^ and reviewed in ^57^), based on key immune similarities between these species ^21,58^. A key limitation of the present studies is that they were not designed to define the protective nuances of a specific vaccine. We did not quantify parameters of *in vivo* protective immunity to pertussis infection such as bacterial titers, nor did we study humoral responses in affected offspring. Future studies utilizing the combined inflammation/vaccine model we now describe could facilitate characterization of these immune perturbations and address their longevity, or possibly identify transgenerational effects, as has been shown in diabetic murine models ^59^. Our present findings also suggest the merit of human studies to determine if infants exposed to antenatal inflammation exhibit altered vaccine-induced immune response patterns, and if so, whether such effects are long-lived. Such knowledge would be important to the optimization of protective immunity in this fragile subpopulation.

## Author Contributions

CMN: contributed to modifications of study design, performed experiments, interpreted data, drafted/edited/revised manuscript, approved final version

DS: contributed to modifications of study design, performed experiments, interpreted data, drafted/edited/revised manuscript, approved final version

JJM: performed experiments, assisted in data analysis, edited/revised manuscript, approved final version

JMK: conceived and designed original study, analyzed and interpreted data, drafted/revised/edited manuscript, approved final version

